## Supplementary figures and legends for "Imaging the response to DNA damage in heterochromatin domains reveals core principles of heterochromatin maintenance"

### Supplementary Figure 1

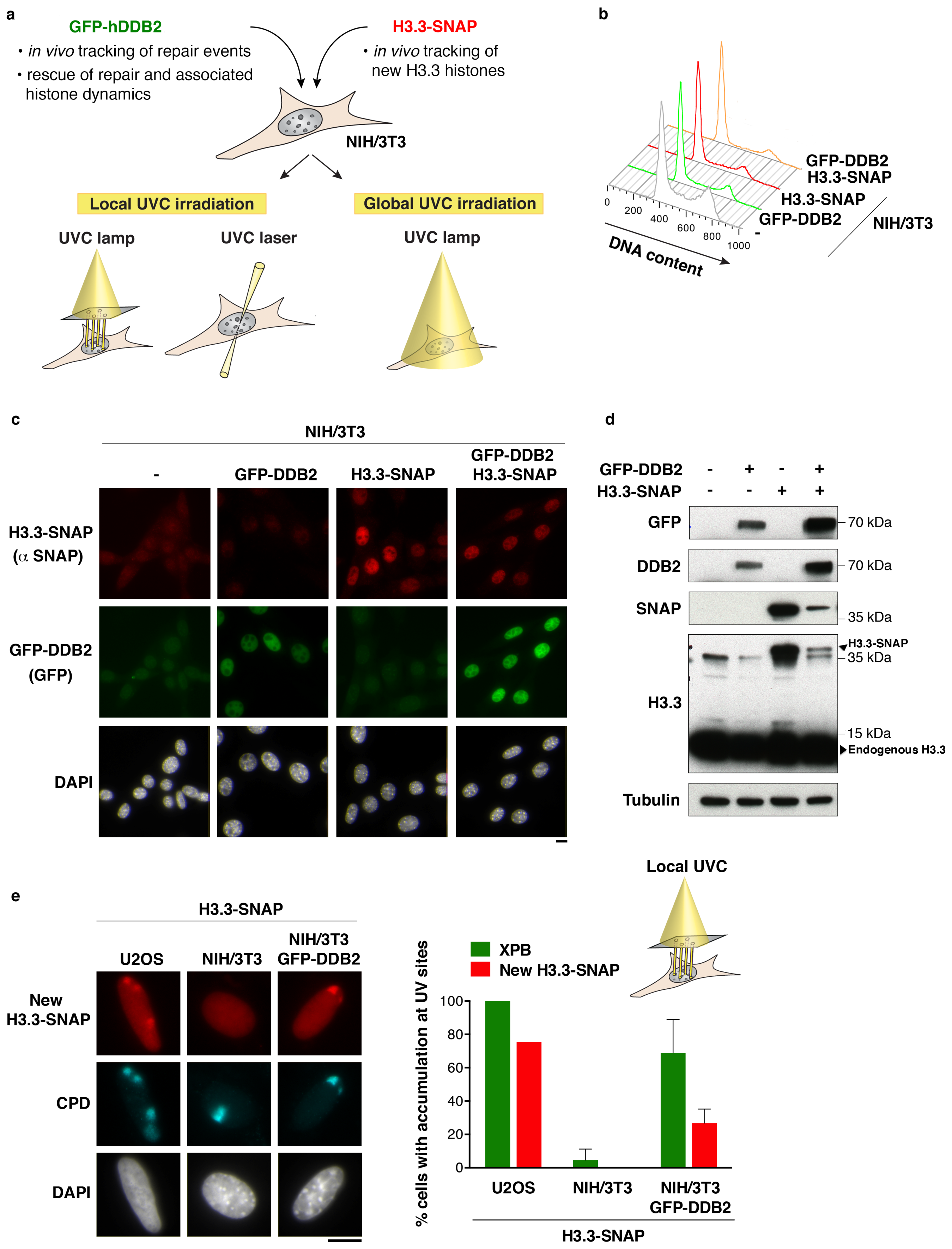

### Supplementary Figure 2

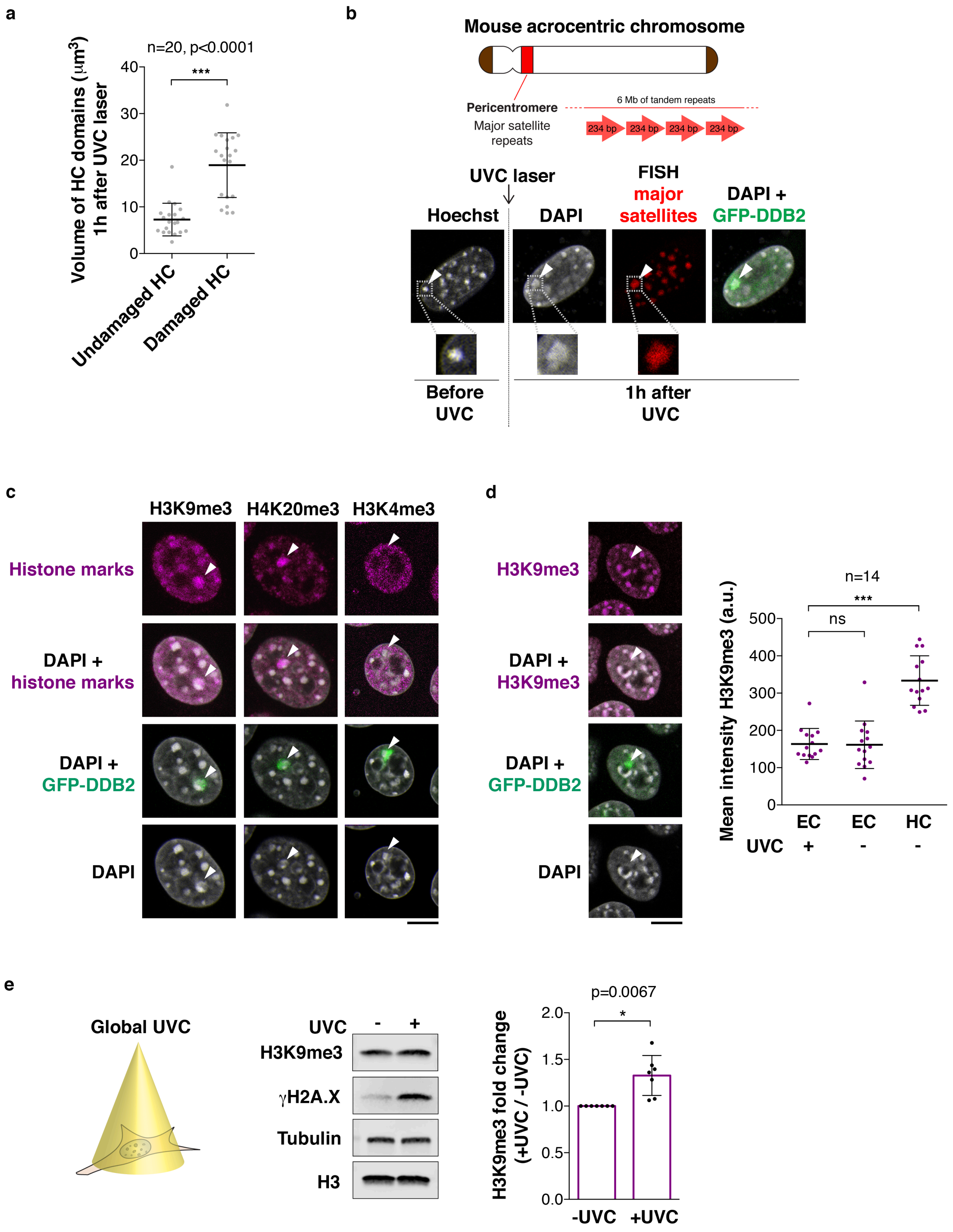

### Supplementary Figure 3

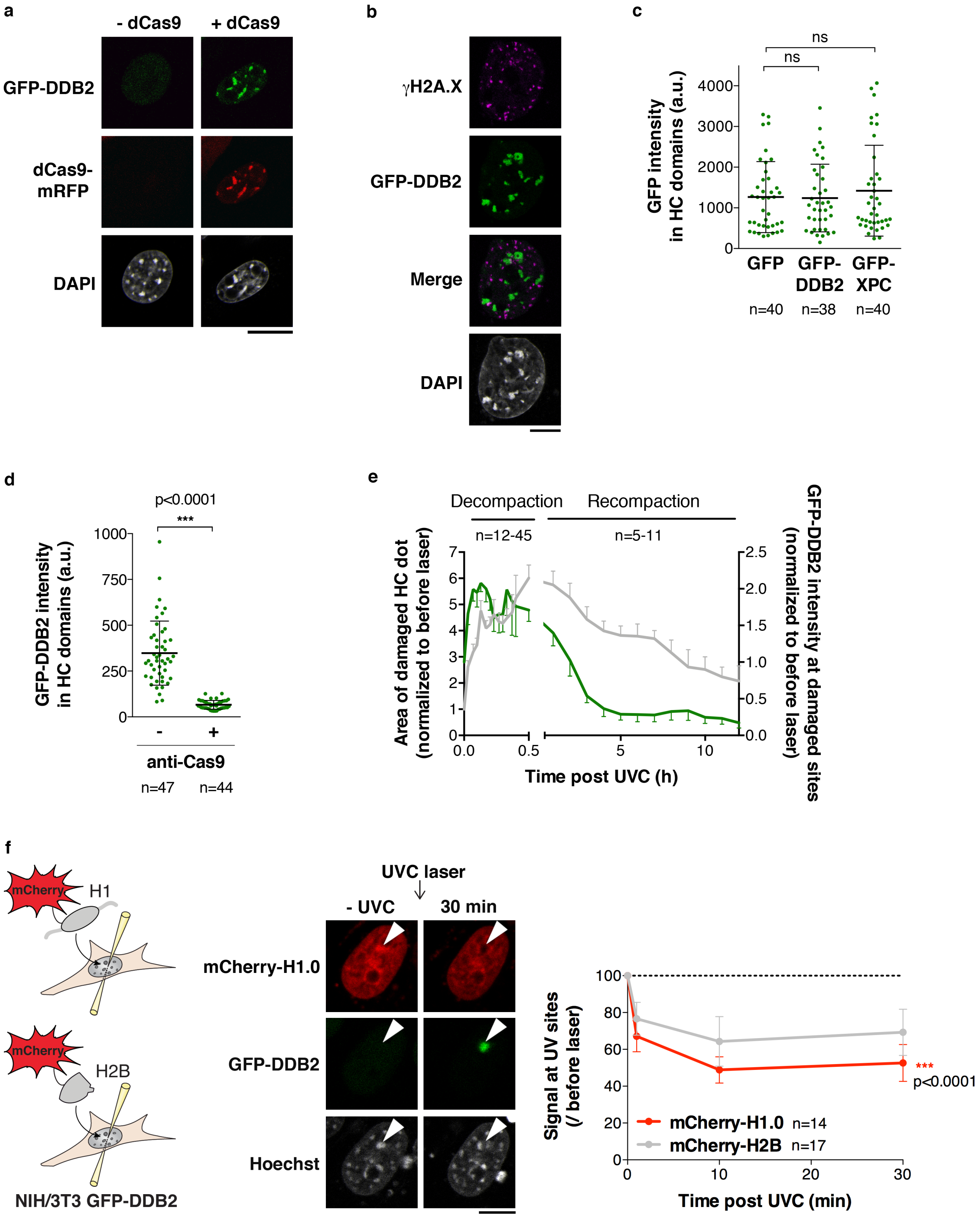

### Supplementary Figure 4

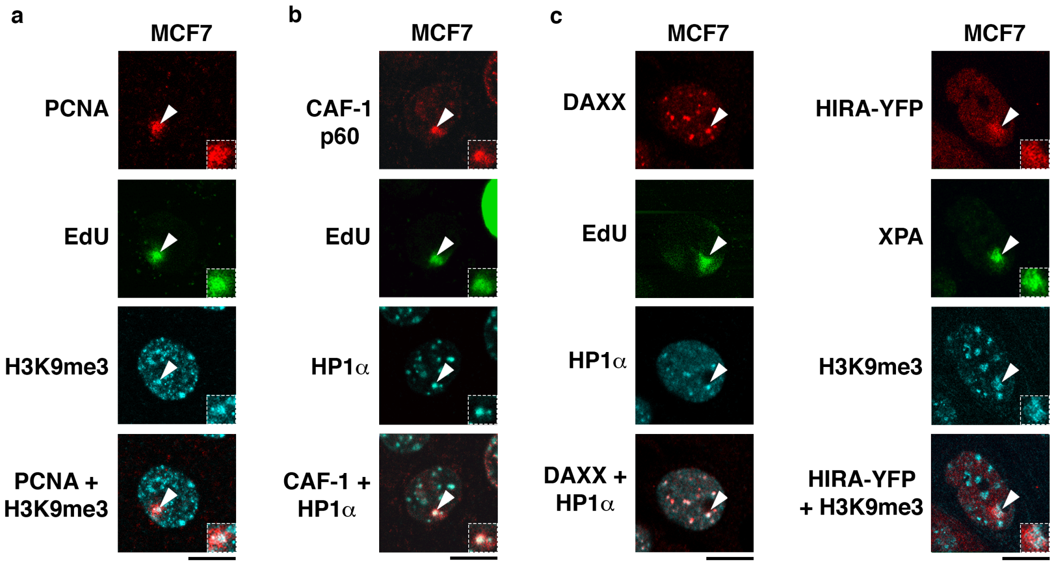

### Supplementary Figure 5

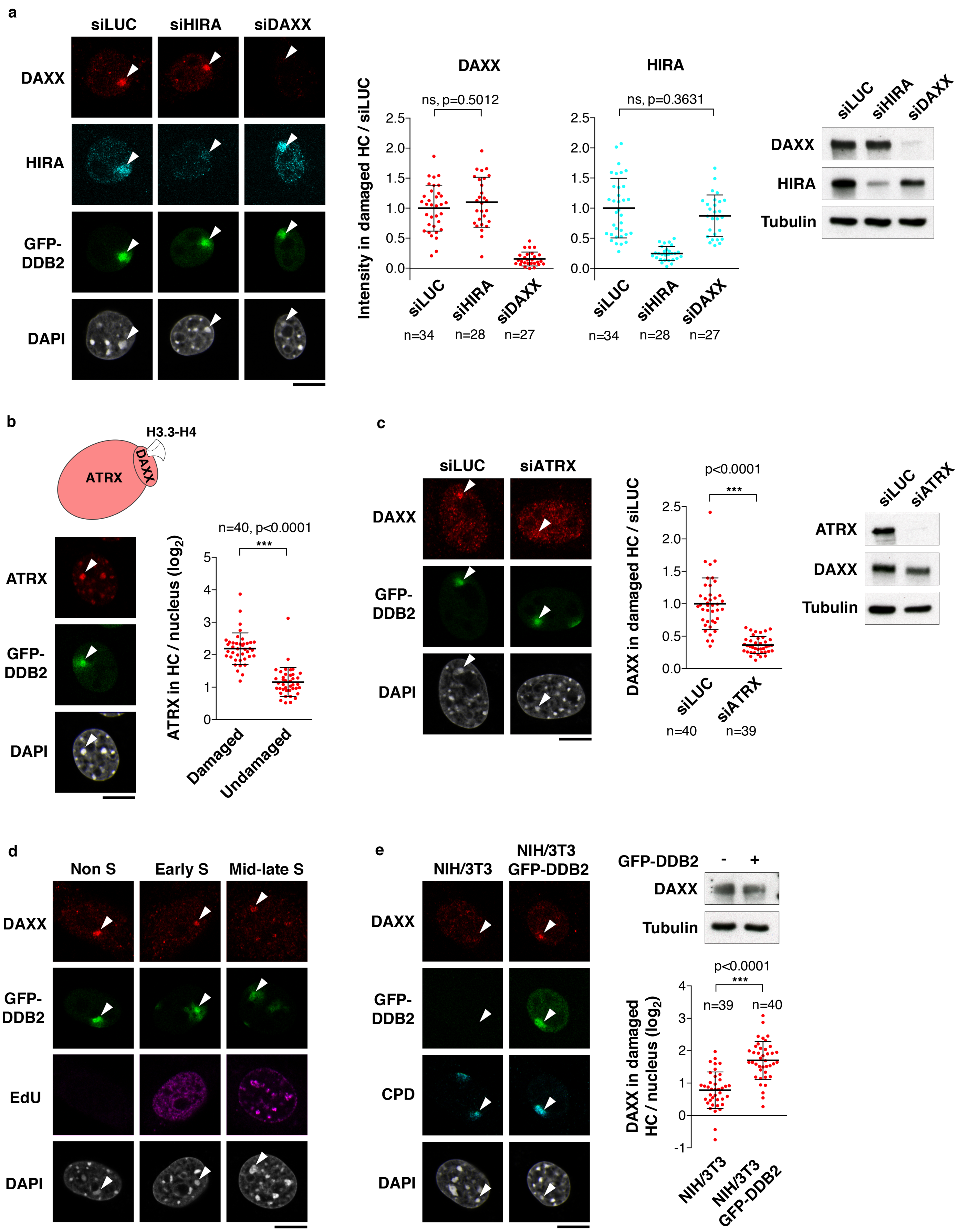

### Supplementary Figure 6

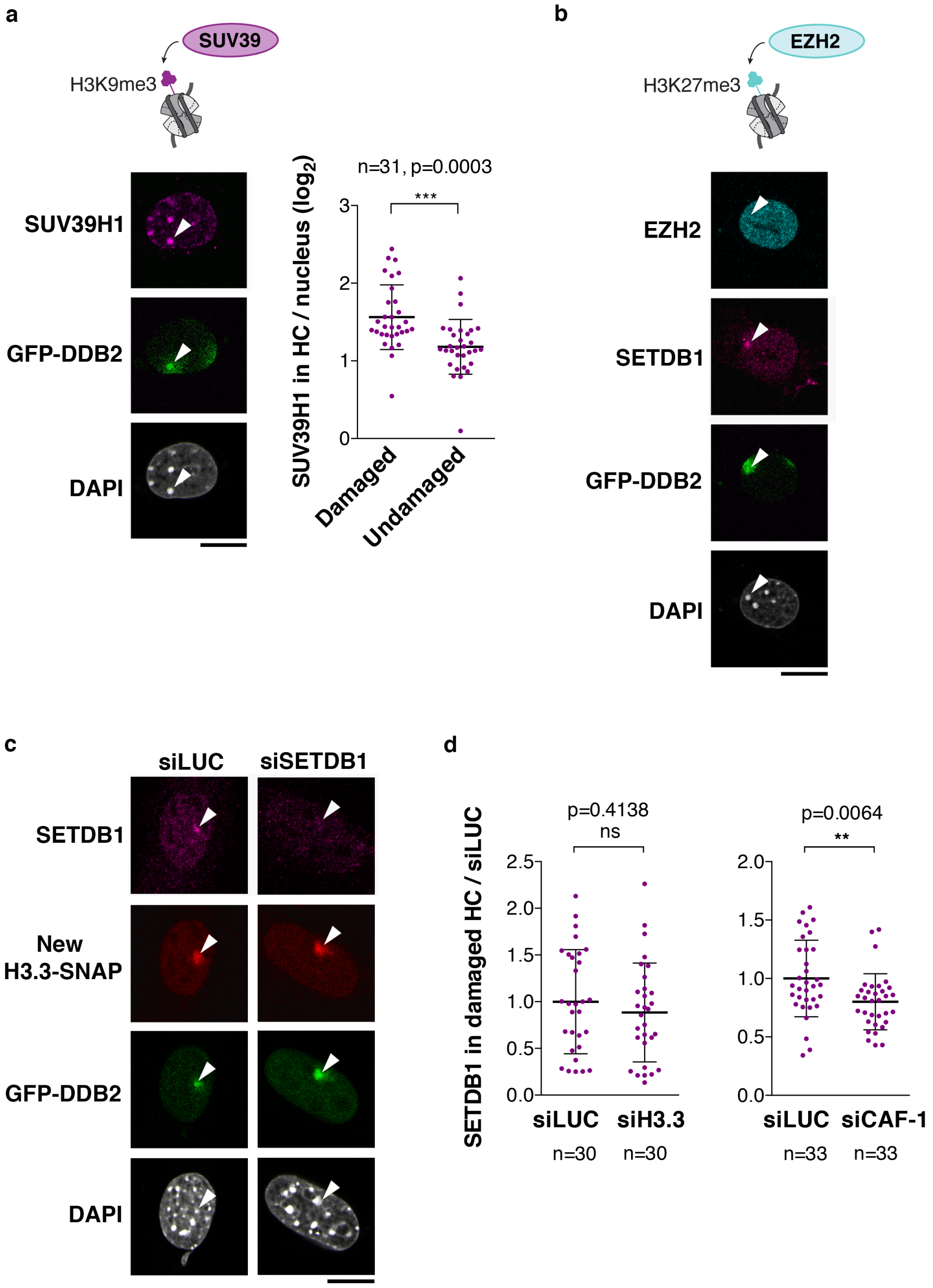

#### SUPPLEMENTARY MATERIAL

**Supplementary Figure 1. Mammalian cellular model for studying heterochromatin maintenance in response to UV damage.**

(**a**) Engineered NIH/3T3 stable cell lines permit to track DNA repair events and to follow H3.3 histone deposition into chromatin in cells exposed to global or local UVC irradiation.

(**b**) Cell cycle distribution analysed by flow cytometry in the NIH/3T3 stable cell lines.

(**c**) H3.3-SNAP and GFP-DDB2 expression analysed by fluorescence microscopy in the indicated cell lines.

(**d**) Total cell extracts of NIH/3T3 stable cell lines (same as in c) analysed by western blot with the indicated antibodies. The top band detected by the H3.3 antibody corresponds to H3.3-SNAP. Tubulin is used as a loading control.

(**e**) New H3.3 histone deposition (red) at UVC damage sites (CPD) analysed by immunofluorescence in the indicated cell lines 45 min after local UVC irradiation through micropore filters. Histograms represent the fraction of cells showing new H3.3 histone accumulation at UV damage sites (red bars). Recruitment of the repair factor XPB to UV damage sites was analysed by immunofluorescence 30 min after local UVC irradiation in the same cell lines and plotted on the same histogram (green bars).

Error bars, s.d. from two independent experiments scoring 150 cells in each experiment. Scale bars, 10 μm.

**Supplementary Figure 2. Decompaction and histone modification changes in UV-damaged pericentric heterochromatin domains.**

(**a**) Decompaction of UVC-damaged chromocenters as compared to undamaged chromocenters 1h after UVC laser micro-irradiation in NIH/3T3 GFP-DDB2 cells. The scatter plot shows the volume of heterochromatin domains based on DAPI staining, measured on reconstructed 3D images (damaged and undamaged heterochromatin domains are from the same nuclei).

(**b**) Schematic representation of major satellite repeats in pericentric regions of mouse chromosomes. Heterochromatin decompaction following UVC laser micro-irradiation is visualized by DNA-FISH of major satellite DNA sequences in NIH/3T3 GFP-DDB2 cells.

(**c**) H3K9me3, H4K20me3 and H3K4me3 in damaged heterochromatin (white arrowheads) analysed by immunofluorescence 1h after UVC laser micro-irradiation in NIH/3T3 GFP-DDB2 cells. H3K9me3 and H4K20me3 are heterochromatin-specific modifications associated with transcriptional silencing while H3K4me3 is a transcriptionally active histone mark used as negative control.

(**d**) H3K9me3 in damaged euchromatin (white arrowheads) analysed as in Fig. 1c. The scatter plot shows H3K9me3 levels measured on reconstructed 3D images in damaged euchromatin domains (EC +UVC) compared to undamaged euchromatin (EC –UVC) and heterochromatin (HC –UVC) in the same nucleus.

(**e**) H3K9me3 levels analysed by western blot 1h30 after global UVC irradiation. Tubulin, loading control; γH2A.X, damage marker. The bar graph represents H3K9me3 abundance in damaged/undamaged conditions.

Error bars, s.d. from seven experiments (e) or from n cells scored in seven (a) or two (d) independent experiments. Scale bars, 10 µm. Zoomed in views of heterochromatin domains (x2.6).

**Supplementary Figure 3. GFP-DDB2 tethering to pericentric heterochromatin domains.**

(**a**) Confocal sections showing the tethering of GFP-DDB2 to pericentric heterochromatin domains of NIH/3T3 GFP-DDB2 cells in the presence of catalytically dead Cas9 (dCas9).

(**b**) Confocal sections showing no overlap between the DNA damage marker γH2A.X and GFP-DDB2 tethered to pericentric heterochromatin.

(**c**, **d**) GFP intensity levels in heterochromatin domains quantified on reconstructed 3D images corresponding to Fig. 2C (c) and Fig. 2D (d).

(**e**) Heterochromatin compaction changes (same data as in Figure 1b) and DDB2 recruitment kinetics upon UVC laser micro-irradiation analysed by live imaging in NIH/3T3 GFP-DDB2 cells stained with Hoechst.

(**f**) Scheme of the experiment for the detection of mCherry-tagged H1 and H2B in live NIH/3T3 GFP-DDB2 cells exposed to UVC laser damage. The levels of H1.0 and H2B are measured in UVC-damaged regions, identified by GFP-DDB2 accumulation (white arrowheads), relative to the whole nucleus at the indicated time points after laser damage. Results normalized to before laser damage are presented on the graphs.

Error bars, s.d. from n cells scored in at least three independent experiments. Scale bars, 10 μm.

**Supplementary Figure 4. Validation in human MCF7 cells.**

(**a-c**) Recruitment of the repair factor PCNA (a) and of the histone chaperones CAF-1 (p60 subunit) (b), DAXX and HIRA (c) to UVC damaged heterochromatin domains (white arrowheads) analysed 30 min (HIRA) or 1h30 (PCNA, CAF-1, DAXX) after local UVC irradiation through micropore filters in MCF7 cells. PCNA, CAF-1 and DAXX are detected by immunofluorescence and HIRA upon transfection of HIRA-YFP. Constitutive heterochromatin is revealed by H3K9me3 or HP1α immunostaining. Sites of UVC damage repair are marked by Ethynyl-deoxyUridine (EdU, repair synthesis) or by immunodetection of the repair factor XPA. Insets show zoomed in views of heterochromatin domains (x1.8). All microscopy images are confocal sections. Scale bars, 10 μm.

**Supplementary Figure 5. DAXX accumulation in UVC-damaged heterochromatin.**

(**a**) Recruitment of DAXX and HIRA chaperones to damaged heterochromatin (white arrowheads) analysed by immunofluorescence 1h30 after local UVC irradiation through micropore filters in NIH/3T3 GFP-DDB2 cells treated with the indicated siRNAs (siLUC, control). siRNA efficiencies are controlled by western blot (Tubulin, loading control).

(**b**) Recruitment of ATRX to damaged heterochromatin (white arrowheads) analysed by immunofluorescence 1h30 after local UVC irradiation in NIH/3T3 GFP-DDB2 cells.

(**c**) DAXX recruitment to damaged heterochromatin upon ATRX knock-down (siLUC, control) 1h30 after local UVC irradiation in NIH/3T3 GFP-DDB2 cells.

(**d**) Recruitment of DAXX to damaged heterochromatin (white arrowheads) 1h30 after local UVC irradiation in NIH/3T3 GFP-DDB2 cells. Cell cycle stages were defined based on staining of replication foci with Ethynyl-deoxyUridine (EdU).

(**e**) Recruitment of DAXX to damaged heterochromatin analysed in the indicated cell lines 1h30 after local UVC irradiation. DAXX total levels are shown on the western blot (Tubulin, loading control).

The scatter plots show DAXX, ATRX and HIRA levels in damaged heterochromatin normalized to the corresponding siLUC experiment (a, c) or log2 fold enrichments compared to the whole nucleus (b, e). Error bars, s.d. from n cells scored in three independent experiments. All microscopy images are confocal sections. Scale bars, 10 µm.

**Supplementary Figure 6. Recruitment of histone methyltransferases to UVC-damaged heterochromatin.**

(**a**, **b**) Recruitment of the histone methyltransferases SUV39H1 (a) and EZH2 (b) to damaged heterochromatin (white arrowheads) analysed by immunofluorescence 1h30 after local UVC irradiation through micropore filters in NIH/3T3 GFP-DDB2 cells. Scatter plots represent log2 fold enrichments compared to the whole nucleus.

(**c**) Accumulation of newly synthesized H3.3 histones in UVC-damaged heterochromatin regions (white arrowheads) upon SETDB1 knockdown (siLUC, control) analysed in NIH/3T3 GFP-DDB2 H3.3-SNAP cells 1h30 after local UVC irradiation through micropore filters.

(**d**) Scatter plots representing log2 fold enrichments of SETDB1 in damaged heterochromatin normalized to the corresponding siLUC experiment upon knock down of H3.3 or CAF-1 (p150 subunit).

Error bars, s.d. from n cells scored in three independent experiments. All microscopy images are confocal sections. Scale bars, 10 µm.

**Supplementary Movie 1: Pericentric heterochromatin decompaction following UVC laser irradiation (22 min kinetics).**

Heterochromatin decompaction visualized by Hoechst staining during the first 22 min following local damage with the UVC laser in a NIH/3T3 GFP-hDDB2 mouse fibroblast nucleus. 12 images were captured at 2 min intervals and are displayed at 2 frames/sec. The resulting motion picture is shown with a superimposed white arrowhead pointing to the laser irradiation site.

**Supplementary Movie 2: Pericentric heterochromatin decompaction and recompaction following UVC laser irradiation (12 h kinetics).**

Heterochromatin decompaction and recompaction are visualized by Hoechst staining during the first 12 h following local damage with the UVC laser in a NIH/3T3 GFP-hDDB2 mouse fibroblast nucleus. 24 images were captured at the following time points: before UVC, 8 min,

30 min, 1h45, and every 30 min till 12 h, and are displayed at 2 frames/sec. The resulting motion picture is shown with a superimposed white arrowhead pointing to the laser irradiation site.
